# Genomic mate-allocation strategies exploiting additive and non-additive genetic effects to maximise total clonal performance in sugarcane

**DOI:** 10.1101/2022.12.19.521119

**Authors:** Seema Yadav, Elizabeth M. Ross, Xianming Wei, Owen Powell, Valentin Hivert, Lee T. Hickey, Felicity Atkin, Emily Deomano, Karen S. Aitken, Kai P. Voss-Fels, Ben J. Hayes

## Abstract

Mate-allocation in breeding programs can improve progeny performance by exploiting non-additive effects. Breeding decisions in the mate-allocation approach are based on predicted progeny merit rather than parental breeding value. This is particularly attractive when non-additive effects are significant, and the best-predicted progeny can be clonally propagated, for example sugarcane. We compared mate-allocation strategies that leverage non-additive and heterozygosity effects to maximise sugarcane clonal performance to schemes that use only additive effects to maximise breeding value. We used phenotypes and genotypes from a population of 2,909 clones phenotyped in Australia’s sugarcane breeding program’s final assessment trials for three traits: tonnes of cane per hectare (TCH), commercial cane sugar (CCS), and fibre. The clones from the last generation of this data set were used as parents to simulate families from all possible crosses (1,225), each with 50 progenies. The breeding and clonal values of progeny were predicted using GBLUP (considering only additive effects) and the e-GBLUP model (incorporating additive, non-additive and heterozygosity effects). Integer linear programming was used to identify the optimal mate-allocation among the selected parents. Compared to the breeding value, the predicted progeny value of allocated crossing pairs based on clonal performance (e-GBLUP) increased by 57%, 12%, and 16% for TCH, CCS, and fibre, respectively. In our study, the mate-allocation strategy exploiting non-additive and heterozygosity effects resulted in better clonal performance. However, there was a noticeable decline in additive gain, particularly for TCH, which might have been caused by the presence of large epistatic effects. When crosses were chosen based on clonal performance for TCH, progenies’ inbreeding coefficients were found significantly lower than for random mating, indicating that utilising non-additive and heterozygosity effects has advantages for controlling inbreeding depression. Therefore, mate-allocation is recommended in clonal crops to improve clonal performance and reduce inbreeding.

## 1. INTRODUCTION

Crop breeding strategies for producing enough food for a growing population have evolved dramatically in different species over the last two decades. Sugarcane breeding programs were initiated from crosses between sugar-rich cultivated species, primarily *Saccharum Officinarum L*., and a wild relative, *Saccharum Spontaneum L*., which provided disease/pest resistance and abiotic tolerance in a range of varieties (Wei and Jackson 2016; Yadav et al. 2020). In modern commercial programs, the emphasis on sugarcane breeding has shifted to crossing highly heterozygous inter-specific hybrids (Wei et al. 2021). The primary objective of this shift is to create genetic variation that can be exploited via selection in the subsequent cycles before reverting to clonal selection. However, the first and most challenging breeding decision is to select the appropriate genotypes as parents for crosses to maximise the performance of progeny for variety development while maintaining genetic diversity in the breeding program (Boeven et al. 2020; Comstock et al. 1949; Technow et al. 2021).

It currently takes 10 to 12 years of field testing to develop sugarcane cultivars (Park et al. 2007). In Australian breeding programs, the majority of parental clones are chosen from advanced-stage trials and commercially grown cultivars, including some from overseas, through variety exchange programs. Parental clones are evaluated by their additive genetic merit for traits including yield, sugar and fibre contents, predicted from BLUP approaches that include pedigree information (Atkin et al. 2009), as well as other characteristics such as disease resistance (Park et al. 2007).

Due to the development of new high-throughput marker technologies, some recent studies have investigated genomic prediction in sugarcane, which permits early selection for complex quantitative traits such as cane yield (Deomano et al. 2020; Gouy et al. 2013; Hayes et al. 2020; Yadav et al. 2021b). Genomic selection (GS) can reduce the breeding cycle by allowing the quick selection of superior genotypes at any stage of the sugarcane breeding program (Voss-Fels et al. 2021). Typically, truncation selection is the first step in GS-assisted breeding programs where high-performing parental clones are selected based on the GEBVs, which are crossed at random to produce the next generation, ensuring that the progenies have a high mean performance (Wei et al. 2021). However, in the presence of non-additive effects, the mean performance of progeny in a clonal breeding program can deviate from the mean breeding value of the parents.

Mate-allocation has been used in animal breeding programs to control inbreeding, maintain genetic diversity, and leverage non-additive genetic effects (Aliloo et al. 2017; Gonzalez-Dieguez et al. 2019; Pryce et al. 2012; Toro and Varona 2010). For clonal breeding programs, mate-allocation strategies are based on the premise that, while selection may be made on GEBVs (heritable effects), the clones employed for commercial reasons should be the results of planned mating that maximise the offspring’s total (clonal) performance. Because clonally propagated crops are primarily polyploid and highly heterozygous, all additive and non-additive genetic effects could be exploited for variety development. Non-additive genetic effects are substantial for a complex trait such as tonnes of cane per hectare (TCH) in sugarcane and can be captured using extended GS models (Yadav et al. 2021b). Furthermore, the presence of these non-additive effects implies a possible gain in overall genetic merit if matings are allocated to exploit dominance (including heterosis) and epistatic effects. In addition, modelling non-additive effects and heterozygosity as effective control of inbreeding would reduce the risk of inbreeding depression in the commercial population, which is substantial for (TCH) in sugarcane (de Azeredo et al. 2016; Silva and Gonçalves 2011). Werner et al. (2020) also advocated that in clonal breeding programs utilising GS, parents should be chosen based on the genomic prediction of cross-performance while considering dominance effects.

We hypothesised that mate-allocation based on predicted progeny performance incorporating non-additive effects would improve the expected performance of the best clones from a sugarcane breeding program. This study aimed to investigate improvements in the prediction of clonal performance in an elite sugarcane population using mate-allocation strategies, where parental clones were selected based on the genomic prediction of cross-performance;

i. Exploiting only additive effects (traditional GBLUP model) to maximise average breeding (additive, GEBV) value, mimicing conventional (random) mating.
ii. Exploiting non-additive genetic effects and heterozygosity in addition to the additive genetic component (extended-GBLUP model for genomic prediction of clonal performance; GPCP) to maximise clonal performance.

We tested this hypothesis with three commercially important traits: tonnes of cane per hectare (TCH), commercial cane sugar (CCS) for sugar content, and fibre content. Stochastic simulations were used to assess the expected clonal performance of selected progeny and inbreeding in the next generation in the above-defined two mate-allocation strategies (i and ii, defined above). Integer linear programming (ILP) was used to determine the mating set that optimises the expected progeny (breeding/clonal) merit.

## 2. MATERIALS AND METHODS

### 2.1 Phenotypes and Genotypes

A 58k SC Affymetrix Axiom cane SNP array was used for genotyping 3,006 elite sugarcane clones from Sugar Research Australia’s breeding program (Aitken et al. 2017). The population used for this study comprised clones evaluated at final assessment trials (FATs) with large plots. From 2013 to 2017, one FAT series was established each year and then harvested over three years (crops) in SRA sugarcane breeding programs in Queensland’s four growing areas of “Northern,” “Burdekin,” “Central,” and “Southern,” with four trials per region and each year. Clones in each series were largely repeated throughout at least three trials within a region, but most were unique to a region. In each trial, 150–300 clones were planted in four-row by 10-m plots using a partially replicated design, with an average replication of 22%. Only the middle two rows were used to collect data, while the outer two rows served as a buffer against competitive effects. More details regarding data acquisition can be obtained from Yadav et al. (2021b).

Before testing the genomic model, the phenotypes were adjusted for experimental and environmental effects across the series, crop, trial, and region to produce the best linear unbiased prediction (BLUPs). The BLUPs were available from commercial partners. There is, however, a possibility that the use of BLUPs as a response variable can result in a double penalty for the genetic effects estimated. However, in the first stage, no pedigree information was included, which indicates that phenotypes did not shrink towards the pedigree before genomic prediction models were fitted. Furthermore, the error variances in the trials (FATs, the final step of the breeding program) were reported to be low due to the high number of replications and large plot size within-region. BLUPs also accommodated small amounts of missing data.

A diploid parameterisation was utilised for genotype calling, with homozygous genotypes for reference and alternative alleles classified as 0 and 2, respectively, and heterozygous genotypes categorised as 1 (Aitken et al. 2016). The SNP array includes almost 58,028 primarily single or low-dosage markers. Aitken et al. (2016) provide detailed information on the array and genotype calling. As quality controls, SNPs with a minor allele frequency of less than 0.01 were excluded, as were SNPs with an Affymetrix QC score of less than 0.6 and at less than 90% of genotypes called across genotyped clones. After quality control, the population included 2,909 clones with 26,086 highly polymorphic SNP genotypes.

### 2.2 Simulation and genomic prediction framework

This work employs two primary streams of data analysis: one for the segregation of target traits in simulated progeny and the other for predicting the breeding and clonal values of those progeny. First, we need marker locations on the genetic map for simulation to generate the recombination process during formation of gametes that are passed from parent to progeny. Of the high-quality SNPs, only 4,502 were mapped on the currently available Q208 sugarcane genetic map. Despite claims of high linkage disequilibrium (LD) levels in sugarcane (Jannoo et al. 1999; Yadav et al. 2021a), identifying genomic areas within the sugarcane genome with genes related to desirable phenotypes requires a large number of markers. Furthermore, a total of 5,920 unpositioned markers on this genetic map were successfully allocated to existing linkage groups using a LD-based algorithm, resulting in a final set of 10,387 SNPs with MAF > 0.01 on the extended genetic map (Yadav et al. 2021a).

#### 2.2.1 Simulation of Phantom progenies

As a first step in predicting the performance of parental crosses, the genotypes of the phantom progeny had to be simulated. To model marker segregation in the progeny population, 70 parents from the overall population were chosen for crossing based on their genomic predicted breeding values (GEBVs). (Fig 1). To account for dioecy, 35 clones were randomly assigned as male and the remaining 35 as female out of 70 parental clones. Following that, all possible crosses (35×35 =1,225) between male and female clones, each with 50 progenies, were simulated by randomly sampling the parental gametes with crossovers using a newly extended map (Yadav et al. 2021a). The progenies were simulated using the R package “SelectionTools,” version 19.3 (http://population-genetics.uni-giessen.de/∼software/), which used Plabsoft software to mimic meiosis using a count-location approach (Maurer et al. 2008). The methodology implies that the average number of crossovers formed on a chromosome is equal to the length of the chromosome in Morgan units.

**Fig 1:**
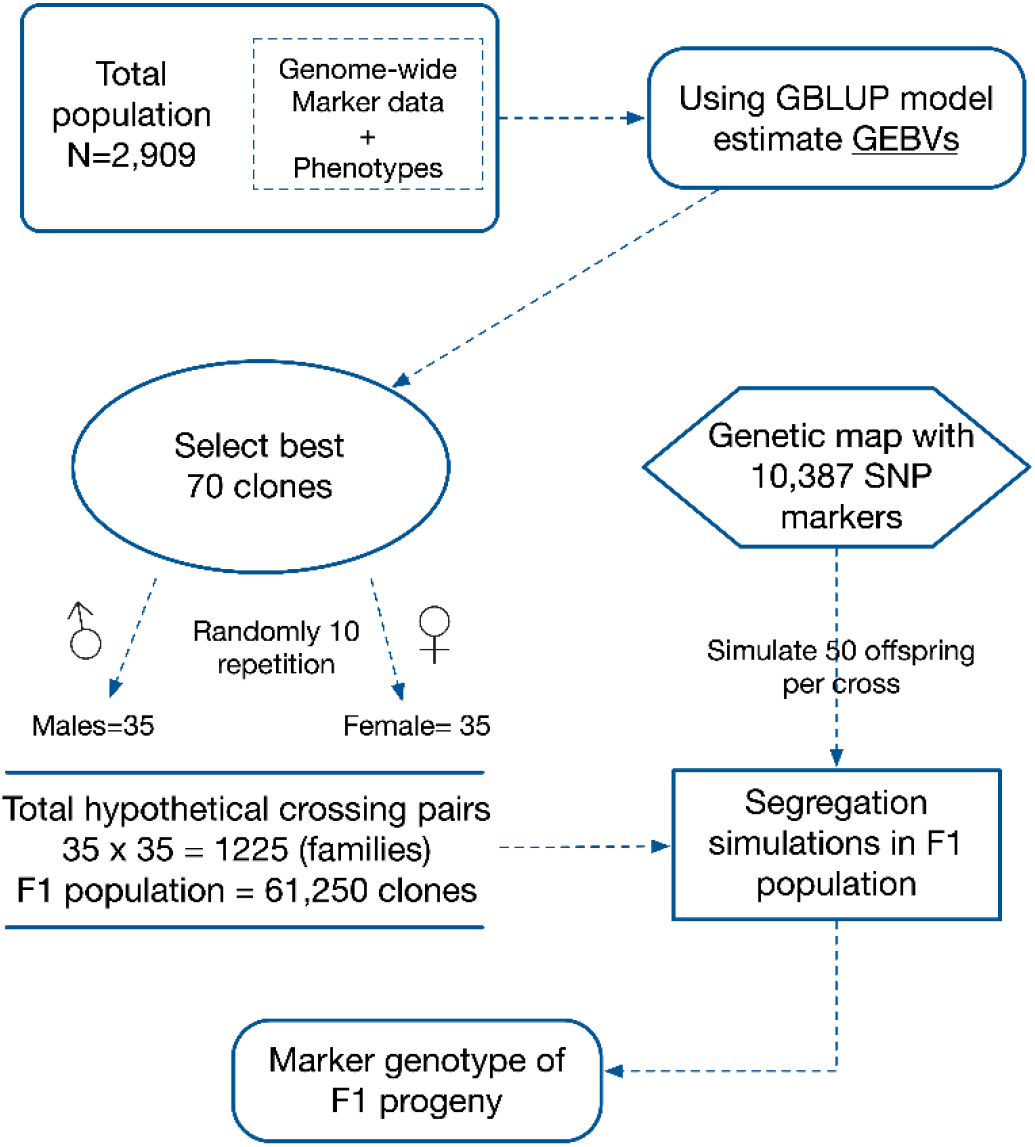
Progeny genotypes: Simulation approach for projecting target trait segregation patterns in phantom progenies from 1,225 families chosen from a pool of 2909 elite clones. For simulation, the data required are genome-wide markers of parental clones and the position of the markers on the linkage map.

Furthermore, the crossover location is distributed independently and uniformly on the chromosome. These assumptions are theoretically equivalent to the underlying Haldane mapping function in the absence of interference (Haldane 1919). In a recent sugarcane study, the same software was used to highlight the different approaches for implementing GS in a simulated breeding environment, taking additive and non-additive effects into account for improving complex sugarcane traits. We conducted ten repetitions in order to eliminate the sampling bias between male and female parents. The parental clones were kept the same in each iteration in order to compare the two strategies for mate allocation.

The additive genetic variance of offspring from a cross can be predicted deterministically using a combination of genome-wide marker effects, a genetic map, and phased parental haplotypes, as shown by Wolfe et al. (2021). Although we have simulated diploid inheritance, which is not appropriate for some regions of the sugarcane genome, however the markers (primarily single or low dosages) in this study were chosen to have diploid like inheritance (Aitken et al. 2016). The limitation of this approach are explored in the discussion.

#### 2.2.2 Prediction of breeding and clonal value of progeny

Single-trait linear mixed models were fitted to estimate genetic variance components for TCH, CCS, and fibre content using the residual maximum likelihood (REML) approach. The variance components were estimated from the entire population of clones, including the parental clones chosen for crossing.

The total population consisted of 64,159 clones, including a training population of 2,909 elite clones with both phenotypes and genotypes. The remaining 61,250 clones (out of 64,159 clones) are simulated progenies generated from 1,225 families from the 70 best parental clones with 50 progeny per family (Fig 2). The performance of these progeny in terms of breeding and clonal value was predicted using a genomic prediction framework based solely on marker profiles (Fig 2).

**Fig 2:**
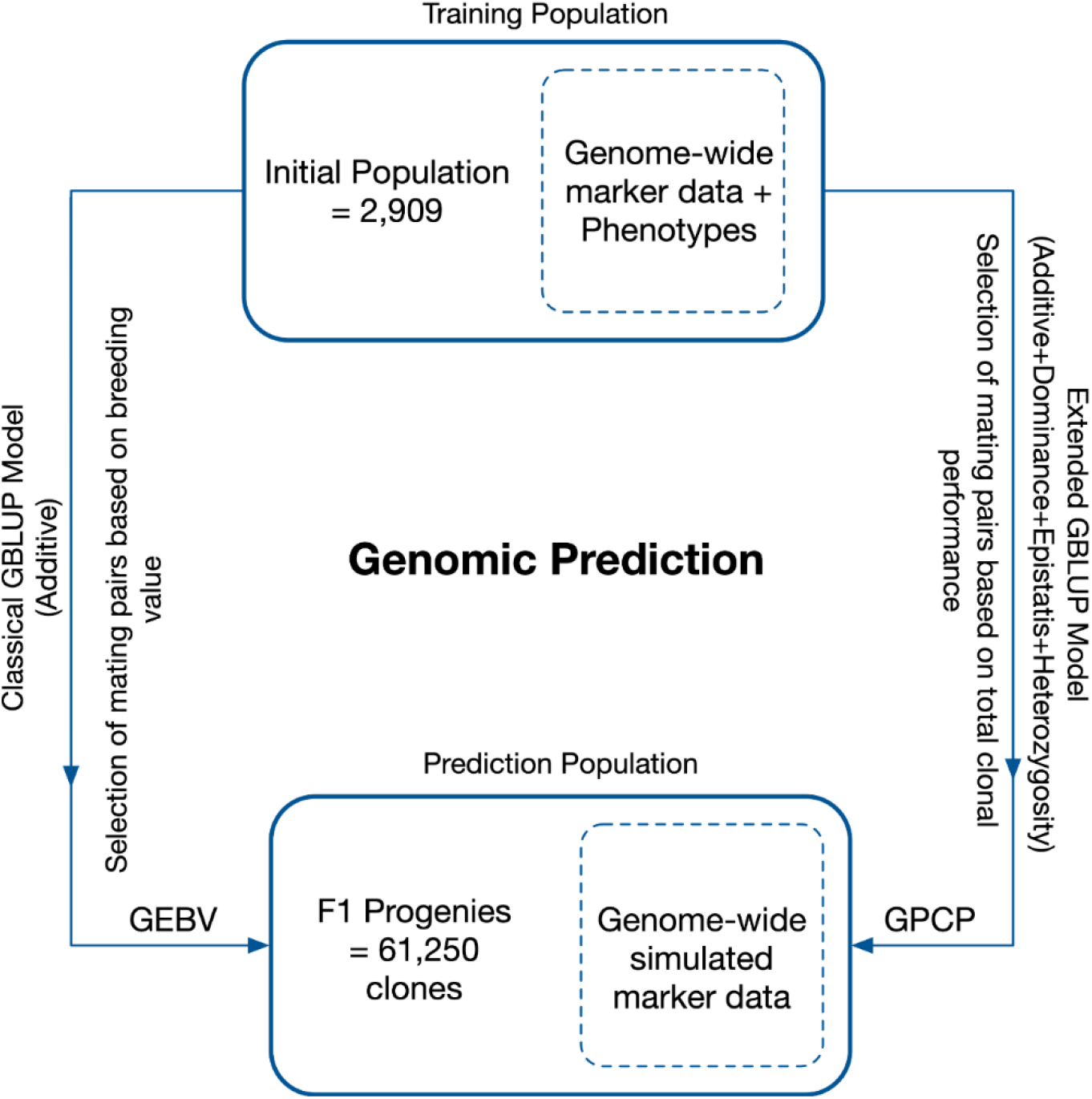
Genomic prediction framework to calculate genomic estimate breeding value (GEBV) and genomic prediction of clonal performance (GPCP) using standard GBLUP and e-GBLUP models, respectively. The model was trained using a breeding population with 2,909 elite sugarcane clones with phenotypes of desired traits and genome-wide markers to predict the 61,250 simulated progenies using their marker profiles only

In matrix notation, the GBLUP and the extended GBLUP model can be represented as follows:

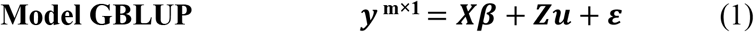

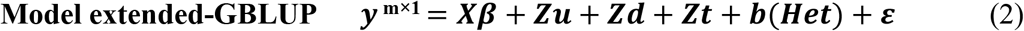

***y*** ^**m×1**^ is a vector of BLUPs for TCH, CCS, or fibre content across the series, crop, trial, and region. ***β***, is a vector of fixed effects, i.e., the overall mean, region, series, trial and crop. ***ε*** is a vector of random residual effects following 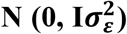, where 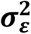 is the residual variance. I is an identity matrix. ***u*** denotes the vector of additive genetic effects. Breeding values (additive effects) were assumed distributed 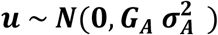, where ***G***_***A***_ is the additive genomic relationship matrix and 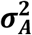, is the additive genetic variance captured by SNP markers. The incidences matrix ***X*** and ***Z*** relate fixed and random effects to observations in ***y*** for a given trait.

The additive genomic relationship matrix, ***G***_***A***_, was calculated according to (Yang et al. 2010) defined as: 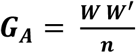, where ***W*** is the incidence matrix of additive genetic effects with dimensions of the number of individuals (**m** = 64,159) and number of SNPs (**n**= 10,387). The elements of *W* are represented by 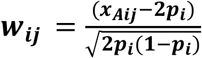, with *p*_*i*_ being the allele frequency at SNP ***i*** (where, *i* =1, 2, …, *n*), and ***x***_***Aij***_ is an indicator variable for additive effects that takes a value of 0, 1, and 2 if the genotype of the ***jth*** individual at SNP ***i*** is *qq, Qq*, or *QQ* (alleles are arbitrarily called *Q* or *q*), respectively. For the GBLUP model (Eq. 1), the mixed model equations (MME) are:

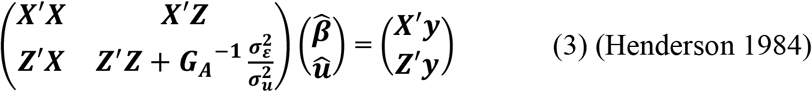

The solution of MME (Eq. 3) is GEBVs which is 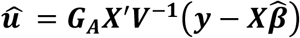, where ***V*** = var(***y***) = var(***Xβ*** + ***Zu*** + ***ε***) = ***ZG***_***A***_***Z***′ + ***R, where R = σ***^**2**^***I***_***m***_, and 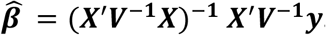. An average of the top 10% of all 50 progenies was taken to determine the expected breeding value for a given cross since we expect only a small percentage of clones will be taken forward for variety development.

In the extended-GBLUP model (Eq. 2), ***d*** and ***t*** are the random vectors of dominance deviation and additive-additive interaction deviation effects. Dominance deviation effects ***d*** were distributed as 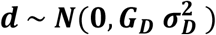, where the genomic relationship matrix for dominance effects was built on genome-wide markers defined by Zhu et al. (2015) as: 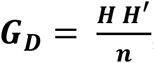, where ***H*** is the incidence matrix of dominance marker covariate matrix with dimensions of the number of individuals (***m***) and number of SNPs (***n***). The elements of *H* are represented by 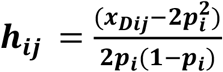, with ***p***_***i***_ being the allele frequency at SNP ***i*** (*i* =1, 2,…, *n*), and ***x***_***Dij***_ is an indicator variable for dominance effects that takes a value of **0, 2*p***_***i***_, and (**4*p***_***i***_ − **2**) if the ***jth*** individual’s genotype at SNP ***i*** is *qq, Qq*, or *QQ*, respectively. This parameterisation of dominance effects ensures orthogonality with additive effects. The additive-additive epistatic deviation effects were assumed to be normally distributed as 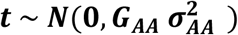. The additive-additive epistatic relationship matrix was represented by ***G***_***AA***_, calculated using the methodology by Vitezica et al. (2017). As a result, the additive-additive genomic relatedness matrix is defined as: 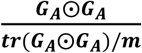 with ***G***_***A***_ ⨀***G***_***A***_ the Hadamard product (i.e., coefficient-wise matrix product) of the additive GRM with itself, ***m*** being the number of individuals and ***tr***, denotes the trace of the matrix (in this case, the trace of the ***G***_***A***_⨀***G***_***A***_ matrix). The corresponding additive-additive interaction variance was represented by 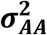, captured using the genome-wide SNP markers. The standardisation by the average of the diagonal elements guarantees that the mean of the diagonal elements of ***G***_***AA***_ is nearly one as it is for ***G***_***A***_ and ***G***_***D***_ resulting in genetic variances estimates on the same scale as the residual variance (Vitezica et al. 2017). Genome-wide heterozygosity (***Het***) was computed from the dominance incidence matrix described in (Yadav et al. 2021b) and fitted in a model (Eq. 2) as a covariate. ***b*** represents the linear regression coefficient of genomic heterozygosity.

The extended-GBLUP model represented by Eq. 2 is theoretically orthogonal to the additive and dominant genetic components. By orthogonality, we imply that there is no covariance between the genetic components; for example, estimates of additive genetic effects remain unbiased even when other genetic components’ effects are present in the model. The additive and dominance GRMs were computed using the GCTA software v.1.93.0b standard algorithm (Yang et al. 2011). The additive-additive GRM was computed from the additive GRM using R v.3.6.2, R-script adopted by Hivert et al. (2021). GRM computation on reasonably large data sets necessitates a substantial amount of memory. GCTA software accelerates the process by building the GRM by block, and each block is allocated to a separate thread, to reduce the computational load. The GRM was constructed as a series of 8 blocks in our implementation. The different variance components of the multiple-GRM models were estimated using Linux-based software MTG2_v2.17 (Lee and Van Der Werf 2016). MTG2 fitted models with the “direct average information” algorithm using REML for variance component estimates.

For the extended-GBLUP model (Eq. 2), BLUPs solutions are:

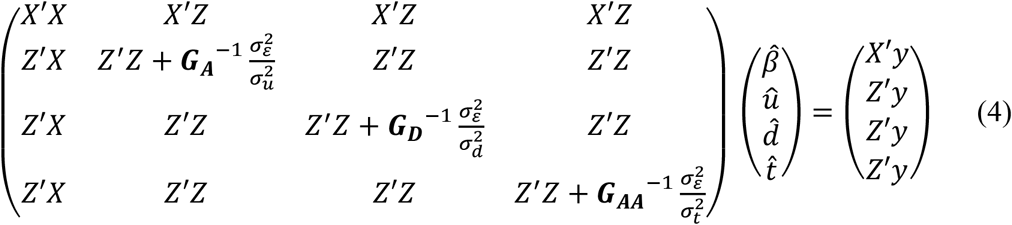

In a mixed model framework, Eq. 4 was used to predict breeding values, dominance deviations, and additive-additive interaction deviation effects. Individual regression coefficients on average heterozygosity (represented by ***b*** in Eq. 2) were computed from the dominant incidence matrix using the approach (Yadav et al. 2021b). The clonal value (*ĝ)* is calculated by summation of predicted random (breeding value, dominance deviations, additive-additive interaction deviation effects, and the genome-wide heterozygosity) effects. The expected clonal value for a particular cross was determined by averaging the top 10% of all 50 progenies.

We used likelihood ratio tests (LRT) to compare the goodness of fit between the nested GBLUP models using ***Test statistic*** = 2 [max(logL(extended GBLUP)) – max(logL(standard GBLUP)], where extended-GBLUP was considered as a complete model. At **p < 0.05**, the test statistic approximates the chi-squared distribution with one degree of freedom. If the ***Test*** 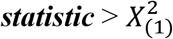 at **p < 0.05**, the model is considered significant.

#### 2.2.3 Inbreeding Coefficients

Inbreeding coefficients of progenies were estimated by the diagonal element of the additive relationship matrix, *G*_*A*_, *whic*h is the genomic relationship of an individual with itself relative to an arbitrary base population (Yang et al. 2010):

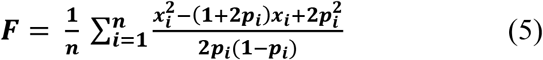

where ***n*** is the total number of SNPs, ***p***_***i***_ being the allele frequency at SNP ***i***, and ***x***_***i***_ is an indicator variable for additive genetic effects and coded as 0, 1, and 2 for *qq, Qq*, and *QQ* genotypes. Furthermore, ***F*** represents the unbiased estimate of the inbreeding coefficient. The mean inbreeding of the progenies from the selected set of mating pairs were computed and averaged over all repetitions to assess the influence of mate-allocation strategies on inbreeding.

### 2.3 Mate-allocation

This study compared two mate-allocation strategies. For the first strategy, the best set of 50 crosses was chosen from all potential mating pairs (1,225) based on their additive value (*û)*, GEBVs to optimise the additive genetic gain in the next generation. In the second strategy, the best 50 crossing pairs were selected based on the predicted clonal performance (*ĝ)*, GPCP to maximise the total genetic values of clones. The reason for choosing 50 crosses was trying to emulate a breeding program that chooses a few varieties with the best clonal performance for commercialization each breeding cycle. The task of optimising additive gain or predicted clonal performance was solved via integer linear programming (ILP) using the R-lpSolve package (http://lpsolve.sourceforge.net/5.5/.) (Berkelaar et al. 2004).

Integer linear programming (ILP) is a static optimisation process used to identify the set of crossing pairs (out of all available crossings) that will maximise progeny performance when the value of one crossing is independent of the value of the other crossing. (Jansen and Wilton 1985). Mathematically, under ILP, a linear objective function is optimised (maximised or minimised) subject to practical real breeding constraints of linear equality and inequality. In our case, the objective function was:

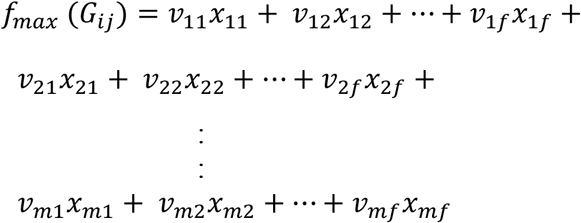

where ***ν***_***ij***_ is the predicted value of the cross (breeding or clonal value) for a particular trait from each selection strategy; ***i*** and ***j*** represent the crossing parents. The practical limitations were that each male parent (***m***) could cross to a maximum of 4 female parents (***f***) and vice versa;

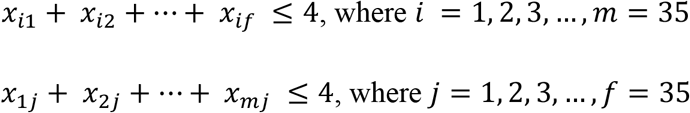

And ***x***_***ij***_, is a binary variable with a value of 0 or 1, where 0 indicates that the crossing between the ***ith*** male and ***jth*** female is not chosen, while 1 represents a selected crossing pair.

In total, 50 crosses were selected. The expected additive gain Δu = *mean*(*û*_50_) − *mean*(*û*_1225_); is defined as the difference between the mean breeding value of selected (50) mating and all possible crossing pairs (1,225). Whereas the expected total genetic superiority Δg = *mean*(*ĝ*_50_) − *mean*(*ĝ*_1225_); is defined as the difference between the total clonal value of selected and all possible crossing pairs. The results of this study are based on an average of ten repetitions of each simulation.

## 3. Results

### 3.1 Variance components and heritabilities

The estimates of variance components, heritabilities, and the maximum log-likelihood ratio values obtained from the two described models (Eq.’s 1 and 2) are shown in Table 1 for the three traits under consideration. TCH had the lowest narrow-sense (*h*^*2*^) heritability estimates (0.18) using the extended-GBLUP (complete) model, compared to CCS (*h*^*2*^ = 0.4) and fibre (*h*^*2*^ = 0.5), which were both relatively high. Estimates of additive genetic variance derived from the GBLUP model differed significantly from those obtained from the extended-GBLUP model, particularly for TCH. For additive variance, the standard errors for TCH using model GBLUP were higher than those in the extended model, whereas the standard errors for the other two traits were the same regardless of the model.

**Table 1.** Estimates of additive and non-additive (dominance and epistatic) variance components, narrow-sense heritability, dominance, additive-additive interaction ratio, log-likelihood ratio (LKH), and heterozygosity effects (*b*) for TCH, CCS and fibre content for 2,909 clones.

| Trait | Model | $\sigma_{\varepsilon}^2$ | $\sigma_A^2$ | $\sigma_D^2$ | $\sigma_E^2$ | $h^2$ | $d^2$ | $E^2$ | LKH | Heterozygosity effects (*b*) |
| --- | --- | --- | --- | --- | --- | --- | --- | --- | --- | --- |
| TCH | GBLUP | 68.51<br>(2.26) | 24.91<br>(3.33) |  |  | 0.27<br>(0.03) |  |  | 7874.67<sup>a</sup> |  |
|  | e-GBLUP | 46.10<br>(3.94) | 15.82<br>(2.9) | 3.08<br>(2.05) | 22.94<br>(4.74) | 0.18<br>(0.03) | 0.04<br>(0.02) | 0.26<br>(0.05) | 7809.28<sup>b</sup> | 92.84*<br>(11.08) |
| CCS | GBLUP | 0.25<br>(0.009) | 0.24<br>(0.02) |  |  | 0.49<br>(0.03) |  |  | 15.05<sup>a</sup> |  |
|  | e-GBLUP | 0.21<br>(0.02) | 0.19<br>(0.02) | ~ 0<br>(NA) | 0.07<br>(0.02) | 0.40<br>(0.03) | ~ 0<br>(NA) | 0.15<br>(0.04) | 25.83<sup>b</sup> | 0.29<sup>ns</sup><br>(0.19) |
| Fibre | GBLUP | 0.81<br>(0.03) | 0.90<br>(0.08) |  |  | 0.53<br>(0.03) |  |  | 1729.24<sup>a</sup> |  |
|  | e-GBLUP | 0.68<br>(0.06) | 0.85<br>(0.08) | ~ 0<br>(NA) | 0.17<br>(0.07) | 0.5<br>(0.03) | ~ 0<br>(NA) | 0.1<br>(0.04) | 1725.26<sup>b</sup> | -0.05<sup>ns</sup><br>(1.22) |

$\sigma_A^2$ = additive genetic variance; $\sigma_D^2$ = dominance genetic variance; $\sigma_E^2$ = epistatic (additive-additive) genetic variance; $\sigma_{\varepsilon}^2$ = residual variance; standard error(SE) in parentheses. LKH=Log Likelihood Ratio. (a-b) models without a common superscript are significantly different at $p < 0.05$ . Model GBLUP= Traditional additive model; Model e-GBLUP = extended-GBLUP includes non-additive effects with average heterozygosity of clones with additive effects. TCH = Tonnes of cane per hectare; CCS (%) = Commercial cane sugar; Fibre (%) = Fibre content. (*b*) = a change in phenotypic mean per 1% increase in heterozygosity for a particular trait. \* means significant effects; ns reflect non-significant effects.

The narrow-sense heritability (*h*^*2*^) estimates for CCS and fibre were comparable in both GBLUP and extended-GBLUP models. When additive, dominance, epistatic genetic effects, and heterozygosity effects were all fitted simultaneously in a model, the estimates of dominance variance for CCS and fibre were nearly equal to zero. In contrast, the dominance effects explained nearly 4% of the phenotypic variance for TCH (Table 1). TCH also had the highest ratio of dominance genetic variation to additive variation (0.19), and additive-additive interaction accounted for around 55% of overall genetic variation. In contrast, additive variance only explains about 38% of genetic variance.

Incorporating non-additive (dominance and additive-additive interaction) genetic effects and heterozygosity effects resulted in a significant reduction in residual variance for all traits compared to the additive model (Table 1, Fig 3). This indicates that a fraction of the non-additive genetic variation was included in the residual variance for the GBLUP model. The additive-additive interaction variation in CCS accounted for about 27 % of the total genetic variance. Fibre exhibited the lowest epistatic variance, accounting for only 17% of overall genetic variance. Based on the log-likelihood, the extended-GBLUP fitted the data better than the regular GBLUP model, regardless of the traits.

**Fig. 3.**
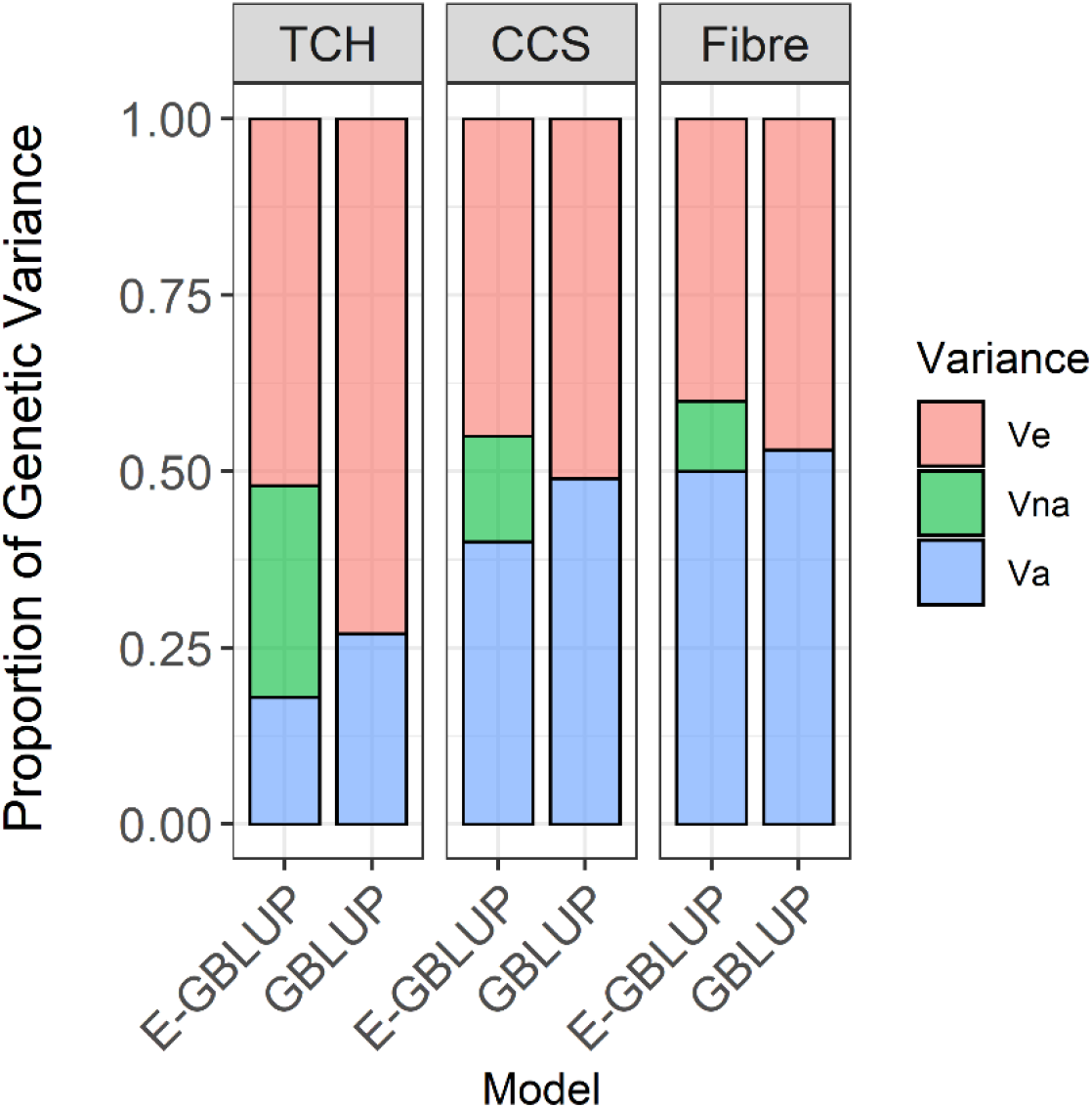
Decomposition of genetic variance into additive and non-additive and residual variance; V_a_ = additive genetic variance; V_na_ = non-additive genetic variance (including dominance and additive-additive interaction variance components); V_e_ = error variance; Model GBLUP= traditional additive model; E-GBLUP = extended GBLUP model by integrating additive, dominance, epistatic and heterozygosity effects.

### 3.2 Effect of heterozygosity

The average heterozygosity across markers was used to determine genome-wide heterozygosity per clone (Table 1). The regression coefficient on heterozygosity for TCH (92.84 ± 11.08) was significant, suggesting an increase in genome-wide heterozygosity is associated with an increase in average cane yield. For CCS and fibre, however, the standard error of the regression coefficient was substantially larger than the heterozygosity estimates and did not differ significantly (Table 1).

### 3.3 Mate-allocation

The total variation of predicted breeding *û* (and clonal *ĝ)* value for all potential crossing pairs (n=1,225), 50 best crosses selected from ILP, and the top decile of best crosses are depicted in Fig. S1 for TCH, CCS and fibre content for one simulation iteration. The mean *û* (or *ĝ)* for all potential crossing pairs (the baseline for our comparisons, n=1,225) was 2.02 ± 0.006 (4.80 ± 0.02) tonnes/ha, 0.24 ± 0.0008 (0.31 ± 0.0009) measured in %, and 0.46 ± 0.002 (0.61 ± 0.002) % for TCH, CCS and fibre, respectively across ten iterations (Table 2). The average expected progeny value of selected matings based on a model that exploited non-additive genetic effects (e-GBLUP) was improved by 57%, 12%, and 16% for TCH, CCS, and fibre, respectively, compared to an additive model (GBLUP) (Table 2).

**Table 2.** Average expected progeny value (EPV_avg_) and genomic measures of average inbreeding of progenies (PIB_avg_) of selected crosses (50) across ten iterations of simulation for TCH, CCS, and fibre with standard deviations in parentheses.

| Mate-allocation | TCH (tonnes/ha) |  | CCS (%) |  | Fibre (%) |  |
| --- | --- | --- | --- | --- | --- | --- |
| Selection of crossing pairs | $EPV_{avg}$ | $PIB_{avg}$ | $EPV_{avg}$ | $PIB_{avg}$ | $EPV_{avg}$ | $PIB_{avg}$ |
| **GEBV<sup>1</sup>** | 5.29<br>(0.06) | 0.04<br>(0.01) | 0.50<br>(0.003) | 0.02<br>(0.01) | 1.16<br>(0.01) | 0.02<br>(0.009) |
| **GPCP<sup>2</sup>** | 8.28<br>(0.07) | -0.046<br>(0.007) | 0.56<br>(0.005) | 0.03<br>(0.01) | 1.34<br>(0.02) | 0.05<br>(0.007) |

<sup>1</sup>Mate-allocation strategy, where crossing pairs would be selected based on GEBVs; <sup>2</sup>Mate-allocation strategy, where crossing pairs would be selected based on GPCPs; GEBV= genomic estimated breeding value; GPCP = genomic prediction of clonal performance exploiting non-additive effects; TCH=tonnes cane per hectare; CCS= commercial cane sugar; Fibre=Fibre content. The standard deviation in parentheses reflects the variation across ten simulation repetitions.

The average genomic inbreeding coefficient of progenies for TCH showed that the selected progenies based on model extended-GBLUP with mate allocation had lower estimates of inbreeding compared to the additive model (Table 2). The negative genomic inbreeding estimates reflect that selected clones in crosses were more heterozygous (less inbred) than the average. The difference between the mean *û* (*or ĝ*) of selected crosses and the mean of all potential matings was called expected additive genetic gain (Δu) and expected total genetic superiority (Δg), respectively. Fig 4 depicts the additive genetic gain (Δu) and total clonal superiority (Δg) obtained with the selected matings are depicted for each mating strategy.

**Fig. 4.**
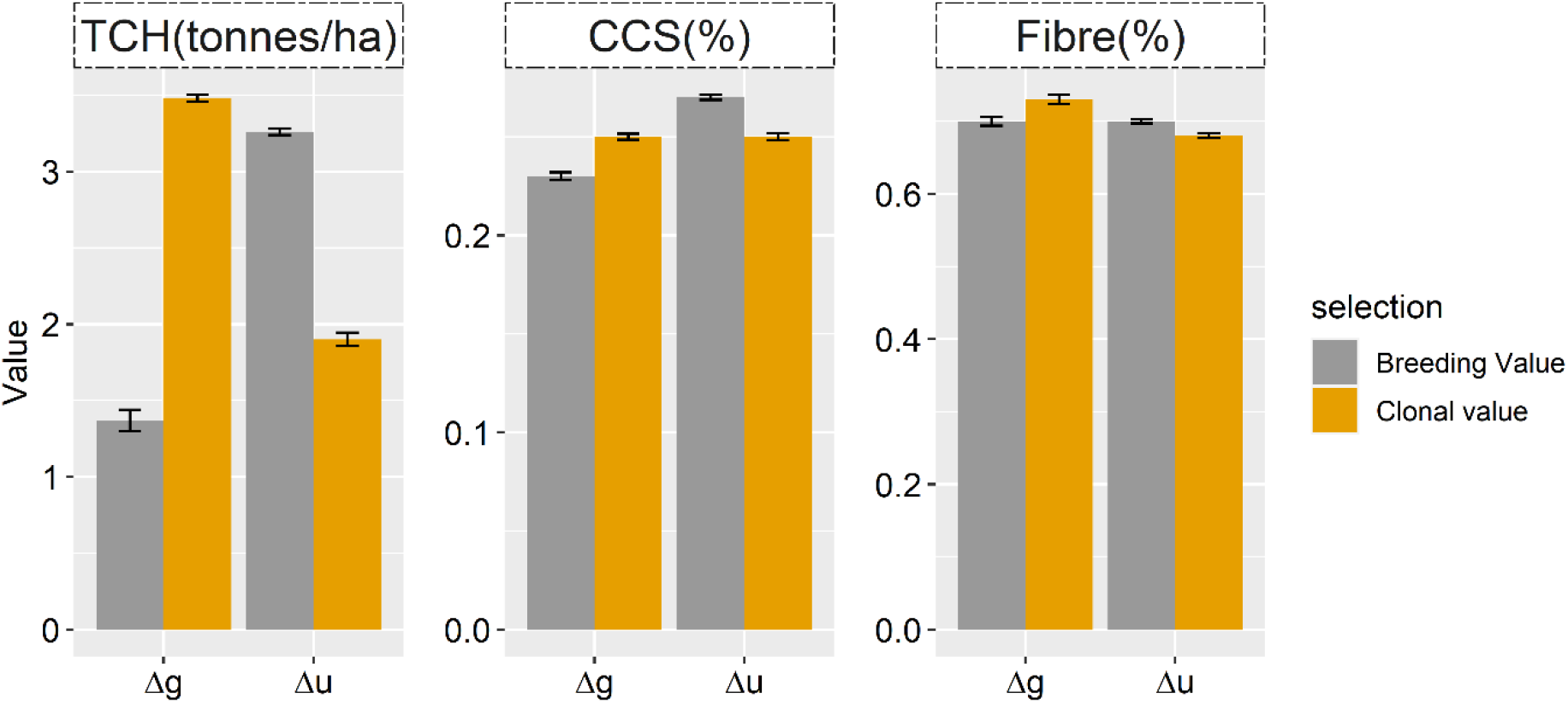
Expected total genetic (Δ*g*) and additive (Δ*u*) gain obtained from 50 selected crosses using a linear programming optimisation algorithm. Δu =*û*_50_ − *û*_1225_; Δg =*ĝ*_50_ − *ĝ*_1225_; where *û*_1225_ (or *ĝ*_1225_) is the mean breeding (or clonal) value of total potential crossing pairs, and *û*_50_ (or *ĝ*_50_) is the mean breeding (or clonal) value of selected crossing pairs using the integer linear programming optimisation technique. The error bar shows the standard error of mean for ten repetitions of the simulation

The expected total genetic superiority of the progeny was higher for all traits when matings were chosen based on clonal performance rather than breeding value, giving the offspring an advantage for TCH, CCS, and fibre, respectively. However, a significant decrease in additive genetic gain was found using the same selection strategy, notably for TCH. No major difference was found between the two strategies in additive (Δu) and expected genetic (Δg) gain for CCS and fibre. The rankings of crossing pairs differed significantly in terms of mate allocation techniques where selection is made on GEBVs or GPCPs. For example, only approximately six crossing pairs out of 50 overlapped between the mate-allocation strategies, for TCH as expected, given its higher epistatic effects. This indicates that different parents were selected to optimise the total additive (or clonal) value.

## 4. Discussion

Our study demonstrates mate-allocation strategies that account for non-additive genetic effects in a sugarcane breeding scheme can improve progeny performance in the next generation. Considering non-additive genetic effects in mating decisions is likely to lead to breeding of higher-performing varieties. The average expected progeny value of selected crossing pairs was improved by 57%, 12%, and 16% for TCH, CCS, and fibre, respectively, when non-additive and heterozygosity effects were exploited. These results aligned with other clonal crops and outbred species (Aliloo et al. 2017; Gonzalez-Dieguez et al. 2019; Werner et al. 2020; Wolfe et al. 2021). In a simulation study, mate-allocation accounting for dominance genetic effects improved offspring performance from 6% to 22% (Toro and Varona 2010). In contrast, Ertl et al. (2014) demonstrated that mate-allocation would provide an estimated overall average genetic superiority in offspring using an empirical data set of cattle, a 14.8% increase in milk yield, and a 27.8% increase in protein yield. Mate-pair allocation in loblolly pine tree breeding (*Pinus taeda L*.) exploiting dominance effects outperformed phenotypic benchmarks by up to 113% (Resende 2014).

As sugarcane breeding programs progress, inbreeding and a reduction of genetic diversity may increase the risk of inbreeding depression. Careful selection and mate-allocation should therefore be performed to balance gain against the negative consequences of increased inbreeding. Lin et al. (2017) also proposed inbreeding controls during mate-allocation when using GS in outbred plants. In this study, selecting crossing pairs while accounting for genome-wide heterozygosity and non-additive effects considerably reduced progeny inbreeding, especially for TCH, and our findings are consistent with previous research (Aliloo et al. 2017; Gonzalez-Dieguez et al. 2019).

A range of mate-selection indices (objective function), such as the optimal contribution (Meuwissen 1997; Wray and Goddard 1994), superior progeny value (Zhong and Jannink 2007), and its extensions, such as optimal population value (Goiffon et al. 2017) and usefulness criterion (Lehermeier et al. 2017; Wolfe et al. 2021), have been developed to balance gains from selection with average inbreeding and co-ancestry. In addition, genetic distance has also been used to evaluate parental genetic differences and, as a result, to predict heterosis and select parents for a cross. However, the relationship between heterosis and genetic distance remains unclear due to inconsistent findings from various studies (Cheres et al. 2000; Liu et al. 2022; Yingbin et al. 2019). However, most clonally propagated crops are polyploid and highly heterozygous, particularly sugarcane. As a result, effectively all gametes produced by any parental genotype are distinct, and all F_1_ descendants are unique (Wei et al. 2021). The overall genetic (additive/non-additive) effects could be used to develop new varieties. Cross-prediction within a population is critical for population improvement in clonally propagated crops. Some recent crop studies also advocated strategies for selecting parents in artificial crosses based on the genomic prediction of cross performance by leveraging the non-additive effects in cereal, e.g. wheat (Lado et al. 2017) and clonal crops, e.g. strawberry and cassava (Werner et al. 2020; Wolfe et al. 2021). Thus, mate-allocation is the most straightforward approach for gaining from non-additive effects among all the strategies.

Our study results revealed a clear relationship between improved progeny performance in the next generation and genome-wide heterozygosity, particularly for TCH, where we observed significant heterozygosity effects. The extend-GBLUP model could result in an overestimation of the dominance variance when heterozygosity is not explicitly taken into account (Iversen et al. 2019). The same observation has been made in our previous study (Yadav et al. 2021b), which used the same data set as this one, when we found a considerable reduction in dominance variance for TCH after including genome-wide heterozygosity per clone in addition to a random dominance term.

Our results show that selecting crossing pairs based on cross-performance exploiting non-additive effects and heterozygosity effects resulted in higher genetic gain (clone performance) than selection based on breeding value, which is consistent with previous studies (Aliloo et al. 2017; Ertl et al. 2014; Fernandez et al. 2021; Resende 2014; Toro and Varona 2010). TCH exhibited the greatest increase in overall genetic gain because non-additive genetic effects account for almost two-thirds of genetic variation. However, our results contrast sharply with mate-allocation techniques in cattle and pig breeding schemes, which claimed a higher predicted total genetic superiority by including non-additive genetic effects to exploit heterosis while reducing expected additive gain by a small or negligible amount (Ertl et al. 2014; Gonzalez-Dieguez et al. 2019). One of the key explanations could be the substantial magnitude of dominance and epistatic effects in sugarcane compared to these studies, which in turn leads to quite different sets of parents being selected to either optimize clonal and additive values in the crossing pairs. ILP was utilized in this study to facilitate the decision-making process in determining the optimal combination of these crossing pairs, taking into account the practical limitations of breeding programs, such as restricting the number of parents that a parent may cross with. This optimisation technique is widely applied in mate allocation research to improve decision-making (Aliloo et al. 2017; Gonzalez-Dieguez et al. 2019; Resende 2014). Although ILPs are NP-hard, it was possible to solve ILP problems using the R-lpSolve package.

Under the Hardy-Weinberg equilibrium condition, the total genetic variance was partitioned into additive, dominance, and epistatic variance. Although the model is theoretically orthogonal, our results demonstrate that additive variance is reduced for all traits when non-additive genetic effects are included in the model. These results were consistent with those obtained when the natural and orthogonal interaction (NOIA) model was applied to the same population of clones where the Hardy-Weinberg equilibrium condition was relaxed (Yadav et al. 2021b). High linkage disequilibrium in modern elite sugarcane clones might explain some confounding effects of additive and non-additive genetic effects (Jannoo et al. 1999; Raboin et al. 2008). Most significantly, a simple diploid model is unlikely to reflect all of the sugarcane’s genomic and genetic complexity. The reduction in residual variance observed in the extended-GBLUP model demonstrates that residual variation contains a significant portion of the non-additive variation in the traditional GBLUP model, which does not account for non-additive effects. Other studies have come up with similar findings (Aliloo et al. 2017; Gonzalez-Dieguez et al. 2019). As a result, the residual genetic variance may include high-order non-additive genetic effects and genetic variation that is not captured by markers, as well as error variance.

To balance long and short-term gain, the sugarcane breeding scheme could be considered to have two (competing) aims: population improvement via recurrent GS focuses on allele substitution effects, which control and increase the frequency of favourable alleles in the population over time, and can be primarily driven for larger genetic gain in the long term; and a variety development pipeline for short-term development makes use of non-additive and heterozygosity effects to improve the phenotypic performance of market-ready clones. Preselection on GEBVs might restrict the opportunity to select alternative clones that might potentially generate progeny with a greater overall genetic value when used in specific matings. It is important to consider that mating, which benefits from non-additive effects, can only increase progeny performance during its implementation and that the benefits resulting from specific combining abilities cannot be accumulated over generations. It is essential to continually update genomic prediction and mate-allocation programs to benefit from non-additive genetic effects in the long run.

Our results are limited to a single population and a single-trait approach; therefore, the proposed approach could select different parental lines for different target traits. In practice, however, expanding the approach to multiple-trait selection is preferable. A simple extension would be to use a selection index that includes multiple traits and then consider the selection index as a new target trait for the existing single trait approach. Another possible modification is directly implementing multi-trait genomic prediction models and evaluating selection lines using an appropriate selection index.

Furthermore, in order to approximate a high-complexity genome, we used simplified assumptions in our simulation schemes. However, genetics in auto-polyploid species is more complicated than in diploid species since more than two alleles may occur at the same locus. As a result, there are additional phases, and recombination and preferential pairing can vary. In addition, there is limited theoretical and experimental information on recombination and segregation in high-ploidy species. And determining which alleles co-occur on each homologous copy gets increasingly challenging as ploidy increases. Furthermore, a reference genome is essential for phasing genotypes in heterozygous polyploids like sugarcane. Unfortunately, the sugarcane community does not have access to the complete reference sequence, which did not allow for leverage of genome-wide phased haplotypes. Nevertheless, the pseudo-diploid markers is a rough approximation of what we used in our study because the majority of the markers in the SNP array are single/low-dose markers. Based on genotype allele count (0, 1 and 2), the software we used to simulate the progenies assumes that the input (marker) data is in the correct gametic phase.

We are aware of our research’s limitations, as well as the fact that the most basic diploid parametrisation (bi-allelic genetic model) is unlikely to fully represent the genomic and meiotic complexity found in sugarcane. This work was inspired by the recent availability of an Affymetrix SNP array to the Australian sugarcane community, which reported 40,000 single-dose SNP, and that single-dose marker can represent more than 75% of polymorphic markers in an individual cross. This simplification also allowed for the use of non-additive genetic effects, which are difficult to comprehend in a high ploidy context. In addition, because of the small number of samples in the training population and environmental variations that obscure the true genotypic values of training samples, this should be considered when breeders select cross combinations. Therefore, conducting field trials to validate our findings would be worthwhile. Further research is required to fully understand the benefits of mate-allocation methods in a larger context.

## 5. Conclusion

Genomic mate-allocation accounting for non-additive genetic effects is a feasible and potentially effective method for improving the clonal performance of future offspring. For our study, mate-allocation strategies that account for non-additive effects were favourable for all traits; progeny performance for cane yield improved by 57% when compared to strategies that solely account for additive effects. Furthermore, when crossing pairs leverage non-additive and heterozygosity effects, the average inbreeding coefficient of progeny was substantially lower, particularly when TCH was the target trait, thus preserving long-term genetic gains. Mate-allocation accounting for non-additive genetic effects and heterozygosity (or inbreeding depression) is easiest to implement compared to all other approaches explained in the literature by directly estimating progeny performance in the next generation. When breeding operations such as establishing a segregating population and the field evaluation of the population requires a significant amount of time and money, systematic planning for selecting crosses becomes even more important. Although genomic mate-allocation improved overall offspring performance, it significantly reduced the expected additive genetic gain, particularly for cane yield. Balancing long-term gains from the selection on additive effects and short-term development of high-performing varieties will be an interesting challenge for future breeding programs.

## Abbreviations

TCH: tonnes of cane per hectare
CCS: commercial cane sugar
Fibre: fibre content
GBLUP: genomic best linear unbiased prediction
e-GBLUP: extended genomic best linear unbiased prediction

## AUTHOR CONTRIBUTION STATEMENT

BJH, KPVF, and SY conceived the study. XW, FA managed field trials and collected and prepared phenotypic data. ED performed a preliminary statistical analysis of phenotypic data. SY conducted simulation and genomic predictions and interpreted the results. VH provided an r-script for the additive-additive relationship matrix. EMR, OP, BJH, KPVF, and LTH provided critical input to interpreting results. SY wrote the manuscript. All authors edited and agreed to the final version of the manuscript.

## SUPPLEMENTAL FIGURE

**Fig.S1.**
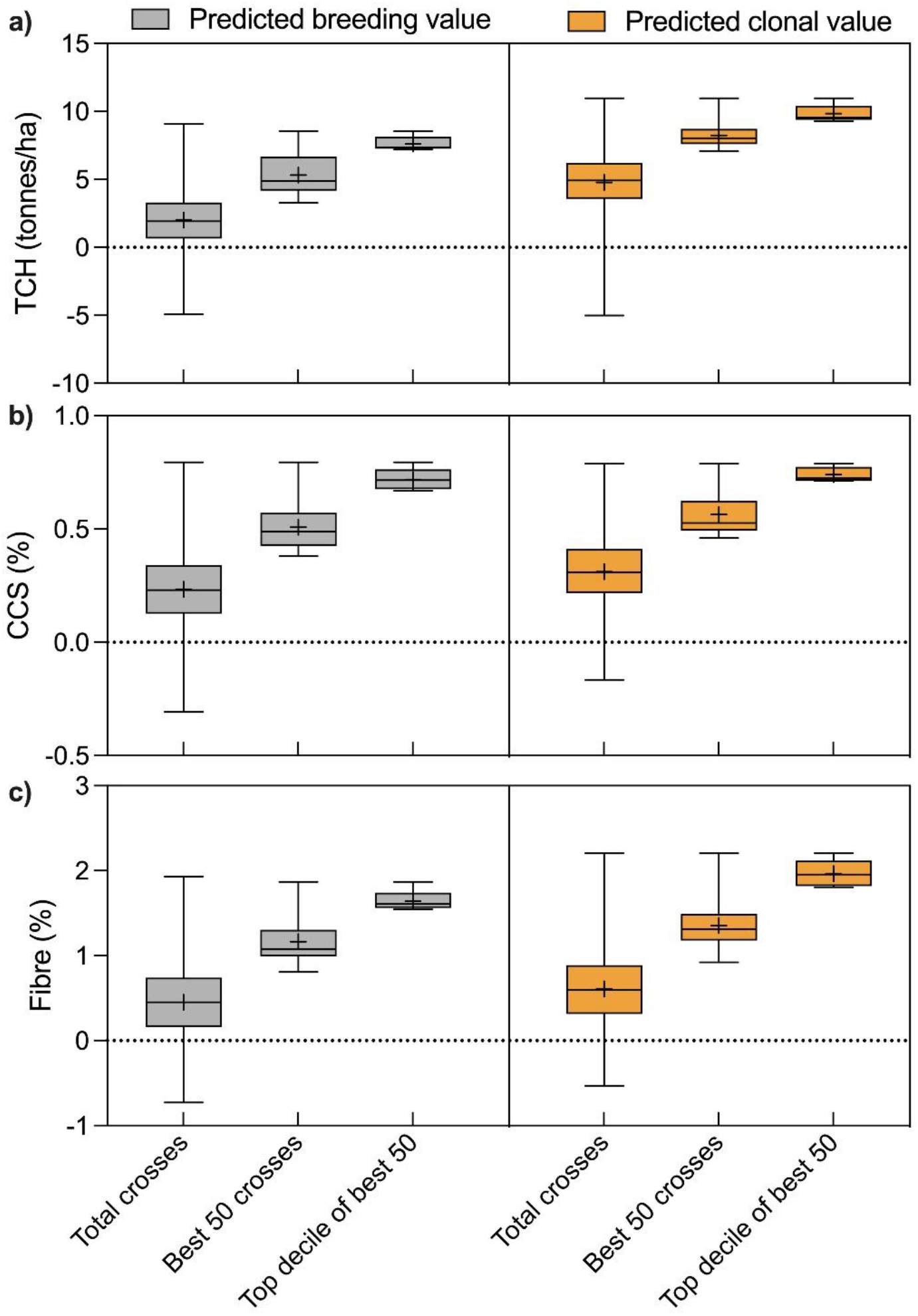
a) Predicted breeding value (left) and clonal value (right) of 1225 crossing pairs, best 50 crosses and top decile of best 50 crosses in one iteration of simulation for TCH; **b)** Predicted progeny (breeding/clonal) value for CCS; **c)** Predicted progeny value for fibre content. The “+” sign represents the mean value, and the solid line across the box represents the median.

